# Neurophysiological Sensitivity to Envelope and Pulse Timing Interaural Time Differences in Cochlear Implanted Rats with Different Hearing Experiences

**DOI:** 10.1101/2025.09.11.675523

**Authors:** S. Fang, T. Fleiner, F. Peng, S. Buchholz, M. Zeeshan, N. Rosskothen-Kuhl, J.W. Schnupp

## Abstract

Cochlear implants (CIs) have successfully restored hearing in more than one million patients with severe to profound hearing loss worldwide. While CIs effectively restore speech perception in quiet environments, sound localization remains challenging for bilateral CI users, particularly their ability to utilize interaural time differences (ITDs). The majority of clinical CI processors use a coding strategy that encodes ITD information only in the envelope of electrical pulse trains rather than their pulse timing, which may contribute to the poorer spatial hearing perception of CI users. We recently demonstrated in a behavioral study on early deafened, bilaterally CI-implanted rats that pulse timing ITDs completely dominate ITD perception, while sensitivity to envelope ITDs is almost negligible in comparison. Building on this, we here investigated the neurophysiological sensitivity of the inferior colliculus (IC) to envelope and pulse timing ITDs at two different pulse rates (900 and 4500 pulses/s) and three different stimulation modulations (5, 20 and 100 Hz) in CI rats with different hearing experiences. Our results indicate that IC neurons exhibit far greater sensitivity to pulse timing ITD than envelope ITD independent of pulse rate, modulation rate or hearing experience. These findings suggest that to improve binaural hearing outcome in bilateral CI users, clinical stimulation strategies should provide informative pulse timing ITDs.

**Graphical abstract:** 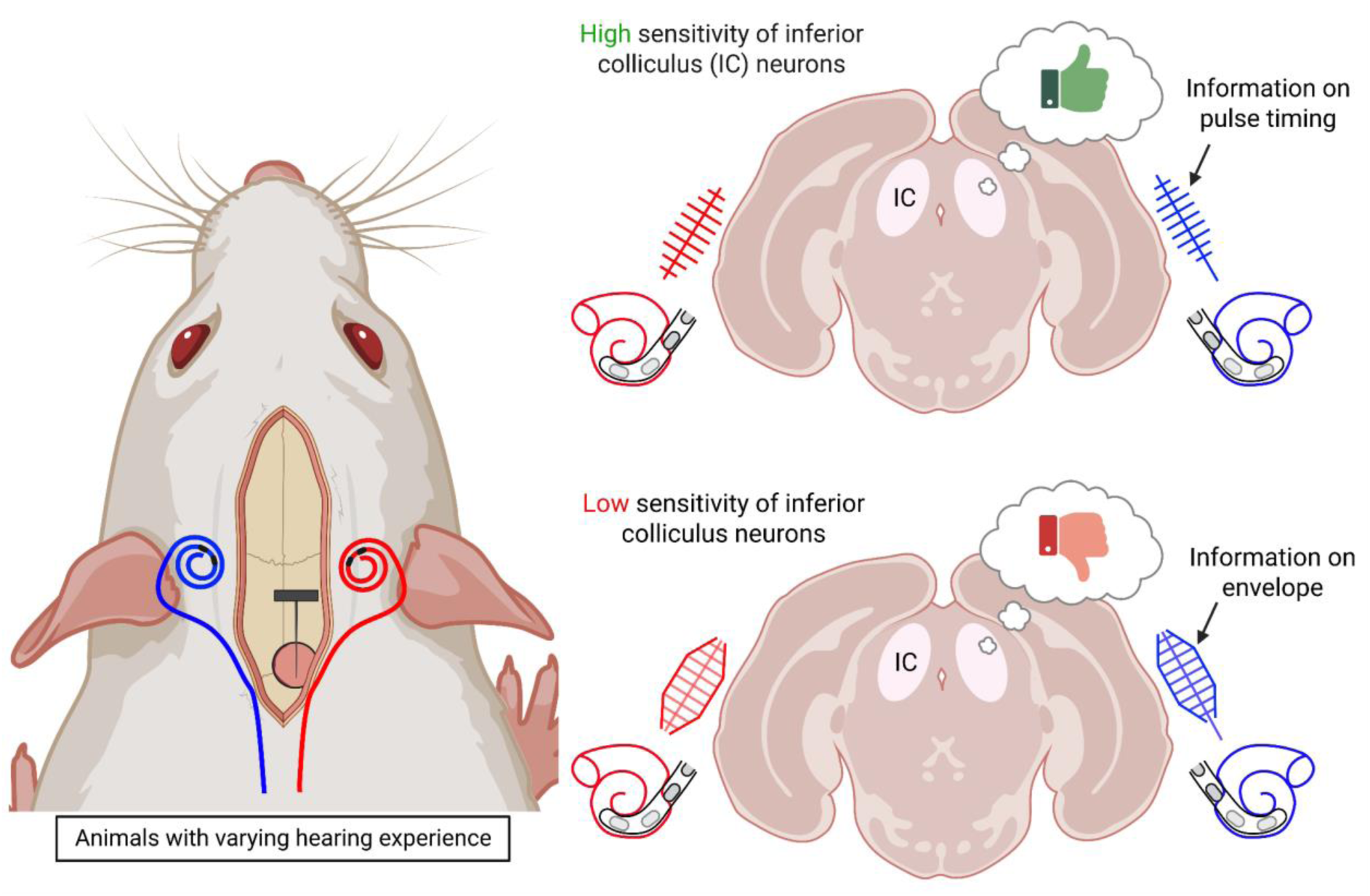

## Introduction

The advent of cochlear implants (CI) has greatly improved the life quality of patients with severe to profound hearing loss by partially restoring auditory sensations, allowing these patients to understand speech and hold conservation. However, the auditory performance of CI patients lags far behind that of normal hearing (NH) humans, especially in hearing tasks requiring high temporal precision, such as the use of interaural time differences (ITDs) for spatial hearing (van Hoesel & Tyler, 2003; Laback *et al*., 2004; Majdak *et al*., 2006; Kan *et al*., 2015). In acoustic hearing, ITD information is conveyed through both the fine structure (FS) and the envelope (ENV) of the acoustic signal. The normally hearing auditory pathway can use sound arrival time differences between the left and right ear as small as 10 to 20 µs to discriminate sound source directions (Klumpp & Eady, 1956; Zwislocki & Feldman, 1956; Laback *et al*., 2007; Thavam & Dietz, 2019). In contrast, ITD thresholds of bilaterally implanted CI (biCI) patients are typically larger than those of normal hearing humans and many prelingually deaf CI patients appear to have no measurable ITD sensitivity at all (Litovsky *et al*., 2010; Laback *et al*., 2015; Ehlers *et al*., 2017; Cleary *et al*., 2022). To be precise, Laback *et al*. (2015) found a mean ITD threshold of around 144 µs testing biCI patients with specialized experimental processors at low pulse rates of around 100 pulses per second (pps; Laback *et al*., 2015). One possible reason for the poorer ITD sensitivity of current biCI patients could be a failure of experience-dependent development of the bilateral auditory pathway due to long-term hearing impairment or the use of unilateral CI, as well as absent early hearing experience (Laback *et al*., 2007, 2015; Litovsky *et al*., 2012; Thakkar *et al*., 2020). However, this is most likely not the main factor for the poorer ITD sensitivity, as adultly deafened CI patients also exhibit increased ITD thresholds (Majdak *et al*., 2006; Laback *et al*., 2007, 2015; Litovsky *et al*., 2012; Thakkar *et al*., 2020; Cleary *et al*., 2022). In addition, recent studies in biCI rats have shown that poor ITD sensitivity is inevitable even in a neonatally deafened (ND) auditory system when they are provided with temporally precise electric pulses directly from the beginning of stimulation (Rosskothen-Kuhl *et al*., 2021; Buck *et al*., 2023). Another possible cause contributing to the poor ITD sensitivity is the design of contemporary CI processors with their very limited ability to provide accurate and precise ITD information via pulse timing (PT) ITD (Wilson *et al*., 1991; Van Hoesel *et al*., 2002; Kan *et al*., 2018). Thus, little to none of the temporal FS of incoming sounds is encoded in the PT of the electrical stimuli. Instead, devices rely largely, or entirely, on encoding the ENV of the sound through pulse amplitude modulation. Consequently, when biCI users receive ITD information, it is typically encoded in the ENVs used to modulate the pulse trains, rather than in the timing of the pulse trains sent to each ear. In fact, commonly used coding strategies such as continuous interleaved sampling (CIS) or CIS derived strategies tend to use pulses fired at a fixed rate independently in each ear, which results in random PT ITDs that may even introduce significant confusion. A notable exception to this is FS4 strategy from MED-EL, which does encode at least some temporal FS information in the pulse timing delivered to low frequency channels.

CIS applies a multiple frequency band-pass filter on the incoming complex sound signal and extracts the envelope information of each frequency band (Wilson *et al*., 1991). A set of fixed rate biphasic pulse trains modulated by extracted ENVs are then sent to the correlated electrode, respectively. Usually, the CI manufacturer shifts the carrier phase of each channel a little to ensure that the adjacent electrode won’t be simultaneously activated, which may help prevent undesirable interactions between adjacent channels. The fixed pulse rate and phase of each channel makes it impossible for contemporary CI operating with CIS based strategies to encode ITD information in the carrier pulse train of the electrical stimulus. Instead, the PT ITDs delivered by these devices will be random numbers, given that either the left or the right ear may be leading by up to half an interpulse interval. It is simply a matter of chance whether, in any given moment, these random PT ITDs agree or disagree with any veridical ENV ITDs that the devices may have been able to encode. Theoretically, it might be possible for the auditory system to “reconstruct” temporal attributes of the signal ENVs at each ear at a higher resolution than the electric pulse sample rate, and to use these to derive ENV ITD estimates independently from the PT ITDs. Whether the auditory pathway of CI patients is capable of doing so in practice is doubtful.

Note that, in the normally developed auditory pathway, cochlear hair cells are known to perform a form of envelope extraction for high frequency stimuli (Palmer & Russell, 1986), but this does not apply to patients with CI, who generally lack normally functioning hair cells, and in whom the auditory nerve is stimulated directly. Consequently, the extensive psychoacoustic literature on ENV ITD sensitivity in normal hearing can be an unreliable guide to what to expect in the CI stimulated case. Furthermore, it is well documented that the auditory nerve has only a very limited dynamic range and resolution for resolving the amplitude of CI stimulus pulses, which is bound to limit the accuracy with which the CI stimulated auditory pathway can reconstruct pulse train ENVs (Zeng *et al*., 1998). Our recent finding that PT ITDs and not ENV ITDs dominate lateralization judgments of neonatally deafened biCI rats trained to lateralize amplitude modulated pulse trains at 900 pps and 4500 pps (Schnupp *et al*., 2025) adds to the suspicion that CI pulse train ENV ITDs may not be represented in the activity of neurons in the auditory pathway of CI users with sufficient accuracy to be a very useful cue for binaural hearing.

In this study, we expand on this behavioral finding and investigate how ENV and PT ITDs are represented by neural discharges in the inferior colliculus (IC), and whether PT ITDs already dominate at this important midbrain auditory relay station. The IC is a well described midbrain structure linked to sound localization and studied neurophysiologically and anatomically for more than five decades (Erulkar, 1959; Oliver & Morest, 1984). The IC is known to integrate and process bilateral auditory information, and many neurons in the IC are sensitive to ITDs (Skottun *et al*., 2001; Hancock *et al*., 2010; Tillein *et al*., 2016; Buck *et al*., 2021). Work of our group investigated the sensitivity of IC neurons in biCI rats to small changes in ITD within the animal’s physiological range, and revealed innate ITD sensitivity in the IC even in the absence of early hearing experience (Buck *et al*., 2021). However, no previous studies have looked at fine-grained, relative ENV and PT ITD sensitivity in IC, specifically over the physiological range, in cohorts of animals with a range of different levels of binaural experience. We therefore conducted a series of neurophysiological extracellular recordings from IC using three different cohorts with different hearing experience: 1) NH rats supplied with acute CIs, tested between P90-150 (Fig. 1, A), 2) ND rats supplied with acute CI, implanted and tested around P90-150 (Fig. 1, B), and 3) ND rats, which were implanted with biCIs in young adulthood, and which had been trained in a stimulus lateralization task with independently varying PT ITD and ENV ITD (Fig. 1, C). Analog multi-unit activity (AMUA) (Kayser *et al*., 2007; Schnupp *et al*., 2015) recorded from these animals was analyzed to compute the proportion of neural response variance explained by stimulus PT or ENV ITD respectively.

**Figure 1.**
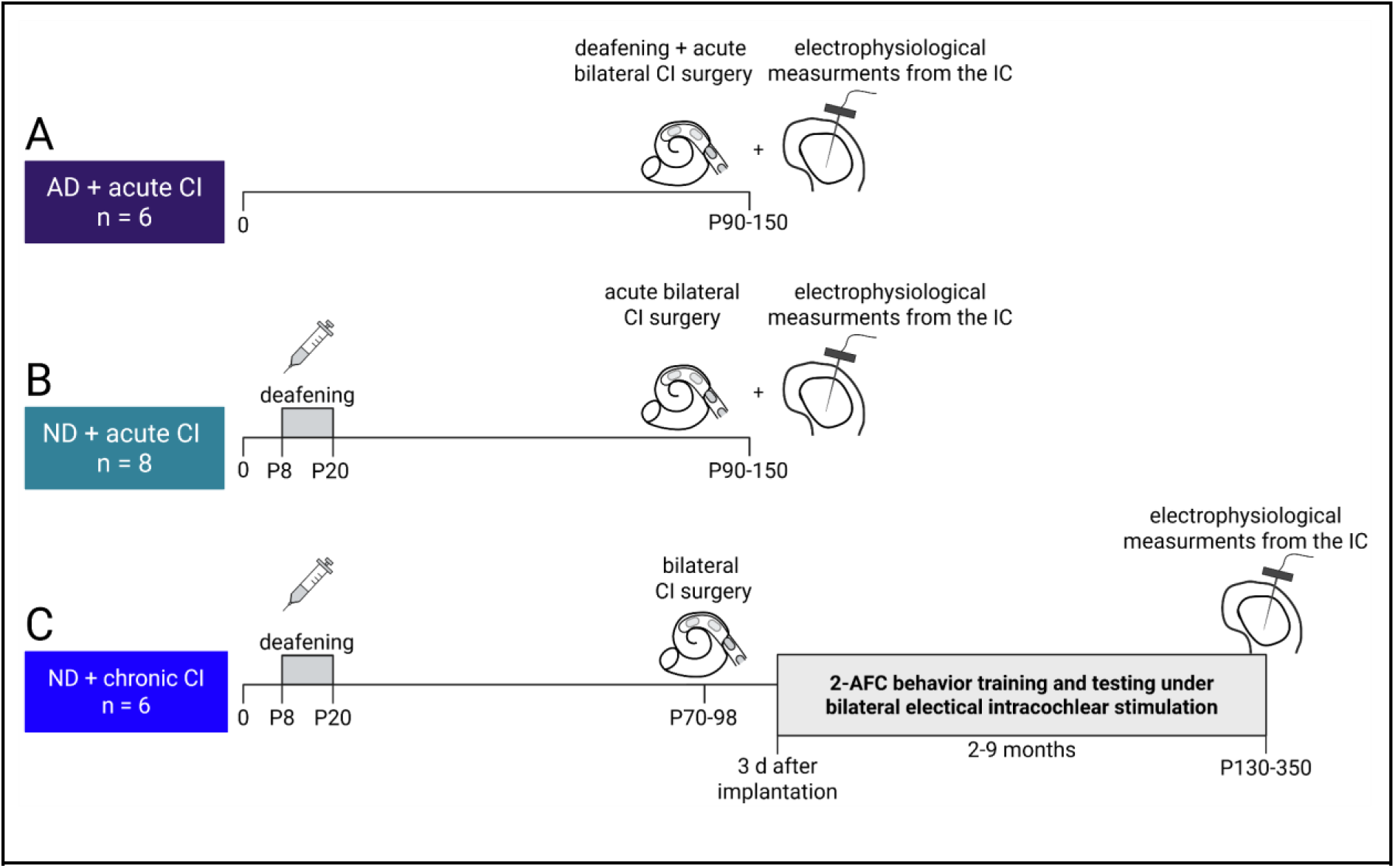
Experimental timetable for the three different cohorts. (A) The first cohort consisted of six animals that were adult deafened (AD) and acutely bilaterally implanted with cochlear implants (CI) immediately prior to neurophysiological recordings from the inferior colliculus (IC). (B) The second cohort consisted of eight neonatally deafened (ND) rats. These rats were acutely supplied with bilateral CI immediately before performing electrophysiological recordings from IC. (C) The third cohort included six ND animals which were bilaterally supplied with CI in young adulthood. From three days after implantation, these animals underwent two to nine months of behavioural training in 2-alternative forced choice tasks (2-AFC) as described in Rosskothen-Kuhl *et al*. (2021). After completion of the behavioral testing, the rats underwent electrophysiological measurements from IC. Illustration created with BioRender.com.

## Material and Methods

All procedures involving experimental animals reported here were approved by the City University of Hong Kong Animal Research Ethics Sub-Committee and conducted under license by the Department of Health of Hong Kong (#16–52 DH/HA and P/8/2/5) and/or the Regierungspräsidium Freiburg (#35-9185.81/G-17/124, #35-9185.81/G-22/067), as well as by the appropriate local ethical review committee. We confirm that all of our methods were performed in accordance with the relevant guidelines and regulations and that our study is reported in accordance with the ARRIVE guidelines. In total, 20 female Wistar rats were obtained (Janvier Labs and Chinese University of Hong Kong Laboratory Animal Service Centre) for this study and underwent terminal electrophysiological recordings. Six animals (AD + acute CI cohort; Fig. 1) were raised with normal hearing, as well as permanent access to water and food until the day of acute bilateral implantation (P90-150) consecutively followed by terminal electrophysiological recording from IC. The remaining 14 rats were neonatally deafened using kanamycin injection protocols (see next section). Of these ND rats, eight remained without electrical hearing experience and with permanent access to water and food until the acute bilateral CI surgery and electrophysiological recording (P90-150; ND + acute CI), while six were bilaterally implanted in young adulthood (P70-98) up to where they had permanent access to water and food and received electrical stimulation while being water deprived (for details see 2-AFC training and testing of the chronic CI cohort) for two to nine months (ND + chronic CI; Fig. 1).

### Deafening and recording of auditory brainstem responses

ND rats were deafened by daily intraperitoneal (i.p.) injections of 400 mg/kg kanamycin from postnatal days 8 to 20 inclusively as described in Rosskothen-Kuhl & Illing (2012) and Rosskothen-Kuhl *et al*. (2018, 2024; Fig. 1). This procedure is known to cause widespread death of inner and outer hair cells (Osako *et al*., 1979; Argence *et al*., 2008). During the injection procedure, Preyer’s acoustic startle reflexes were tested daily and only observed at the early injection period (∼ P14-16; Jero *et al*., 2001). The deafening effect was confirmed by testing the auditory brainstem response (ABR) to broadband click stimuli and pure tones before the surgery. For a subset of the animals, the absence of hair cells in the cochlea was also histologically confirmed.

For ABR recordings, all animals were anesthetized using ketamine (80 mg/kg), and xylazine (12 mg/kg). Acoustic stimuli were delivered to each ear separately through hollow ear bars. Click stimuli consisted of 500 μs white noise pulses presented at 23 Hz, with 400 presentations per intensity level from 0-95 dB sound pressure level (SPL), in 10 dB steps. Pure tone stimuli were presented at frequencies of 3, 6, 9, and 12 kHz, with intensities ranging from 30-100 dB SPL. For details see Rosskothen-Kuhl *et al*. (2021). Rats of the AD + acute CI cohort were not systemically deafened prior to implantation and recording. Instead, conductive hearing loss was induced during the surgical approach to the cochlea, which required removal of the tympanic membrane and ossicular chain to open the cochlea via cochleostomy (Fig. 1).

### Cochlear implantation and measurement of electrical ABR

Rats of the two acutely implanted cohorts (AD + acute CI and ND + acute CI) were bilaterally supplied with CI directly prior to in vivo electrophysiological recordings. In contrast, animals of the chronically implanted cohort (ND + chronic CI) were bilaterally supplied with CI in young adulthood (P70-98) followed by two to nine months of 2-alternative forced choice (2-AFC) behavioral training and testing as described previously in Rosskothen-Kuhl *et al*. (2021), Buck *et al*. (2023), Buchholz *et al*. (2024), Schnupp *et al*. (2025) and briefly below (Fig. 1).

CI implantation was performed as described in previous studies of our group Rosskothen *et al*. (2008), Rosskothen-Kuhl *et al*. (2021), Buck *et al*. (2023). Briefly, rats were anesthetized using ketamine (80 mg/kg) and xylazine (12 mg/kg). During the surgical and experimental procedures, the body temperature was maintained at 38°C using a feedback-controlled heating pad. The vital signs and depth of anesthesia of the animals were frequently checked. The animal was first positioned with the external auditory canal facing upward, and a retroauricular approach was used to expose the tympanic membrane and the auditory tuberosity. The tympanic membrane was punctured, followed by removal of the malleus and incus, exposing the cochlea. The bullar wall around the cochlea was removed using a rongeur to widen the field of view. Finally, a 0.6 mm diameter drill was used to perform a cochleostomy over the middle turn. CI electrodes with two electrode sites, one serving as stimulation site, the other as ground, were inserted through the cochleostomy into the middle turn of the cochlea corresponding to the ∼8 kHz region. In the ND + chronic CI cohort, CI arrays were connected to a custom head connector as previously described in Buck *et al*. (2023), Buchholz *et al*. (2024) and Schnupp *et al*. (2025). The head connectors were screwed onto the skull and fixed with dental cement. In order to get access to the area directly above the IC for electrophysiological recordings, an 8 % agar block was placed on the skull. All implanted CI animal arrays were obtained from MED-EL Medical Electronics, Innsbruck, Austria.

After CI implantation, as well as directly before recordings in the ND + chronic cohort, electrically evoked ABRs (EABRs) were measured in all cohorts, as described in Rosskothen-Kuhl *et al*. (2021), to confirm the effectiveness of bilateral CI stimulation. The EABR amplitudes corresponding to different current intensities were used to initially estimate the minimally effective electrical stimulus amplitude.

### 2-AFC training and testing of the chronic CI cohort

Prior to the terminal electrophysiology experiments described here, the six animals of the ND + chronic CI cohort had undergone two to nine months of training and testing in 2-AFC psychoacoustic ITD and ILD lateralization tasks. For these tasks, the animals were placed in the cage of a custom-made behavioral setup with three water spouts on one side. Licking the central water spout triggered binaural pulse trains very similar to the ones used in the electrophysiological experiments described here. The animals then had to choose one of the two lateral water spouts as indicated by the stimulus ITD to obtain a water reward. Incorrect responses triggered a timeout of a few seconds. All six chronic CI rats of this study developed good behavioral ITD and ILD sensitivity as shown in Buchholz *et al*. (2024).

### Extracellular multi-unit recording

Rats were anesthetized using ketamine (80 mg/kg) and xylazine (12 mg/kg). For maintenance of anesthesia during electrophysiological recordings, a pump delivered an i.p. infusion of 0.9% saline solution of ketamine (17.8 mg/kg/h) and xylazine (2.7 mg/kg/h) at a rate of 3.0 ml/h in the ND + chronic CI cohort. In both AD + acute CI and ND + acute CI cohort, the level of anesthesia was assessed by pedal reflex (firm toe pinch) every 20 minutes, and anesthetic delivery was adjusted as appropriate to maintain surgical plane of anesthesia. All electrophysiological recordings were terminal and after the last recording, rats were euthanized using overdosage of ketamine and xylazine (ND + chronic CI cohort), or overdosage of Dorminal 20% in AD + acute CI and ND + acute CI cohort. To access the IC, rats were placed in a stereotaxic frame and a craniotomy was performed just anterior to lambda and just lateral to the midline suture to expose the occipital cortex overlying the IC. Recordings were mainly obtained from the left IC in AD + acute CI (with 3 penetrations from right IC) and ND + acute CI cohorts, while all recordings of the ND + chronic CI cohort were from the right IC. A single-shaft silicon electrode array (ATLAS Neuroengineering, E32-50-S1-L10) was inserted dorsoventrally into the IC. The electrode comprised 32 platinum recording sites spaced at 50 µm intervals. For each penetration, the tip of our electrode was initially inserted to a depth of ∼4.5 mm from the brain surface. We then searched for responses by increasing the electrode depth in small increments while presenting isolated stimulus pulses at a rate of 1 to 2 per second and ranging in intensity from 0 to 12 dB above 100 μA, depending on the animal’s EABR response. The search stopped when clear responses were observable as acoustic and visual signals on the majority of recording sites. In no case was it necessary to penetrate more than 0.5 mm below the initial 4.5 mm depth. Neural signals were amplified and digitized with an INTAN headstage connected via a PZ5 digitizer by Tucker-Davis Technology (TDT Alachua, FL) to a RZ2 Bioamp processor (TDT) to be sampled and recorded at a sample rate of 24.414 kHz.

### Stimulus design

The stimulation current pulses utilized in this study were generated by a TDT IZ2MH programmable constant current stimulator with a sample rate of 24.414 kHz.

The electrical stimuli were pulse trains (900 pps or 4500 pps) modulated by a Hanning window. The modulation frequencies of 5 Hz, 20 Hz, and 100 Hz resulted in stimulus durations of 10 ms, 50 ms, and 200 ms (Fig. 2).

**Figure 2.**
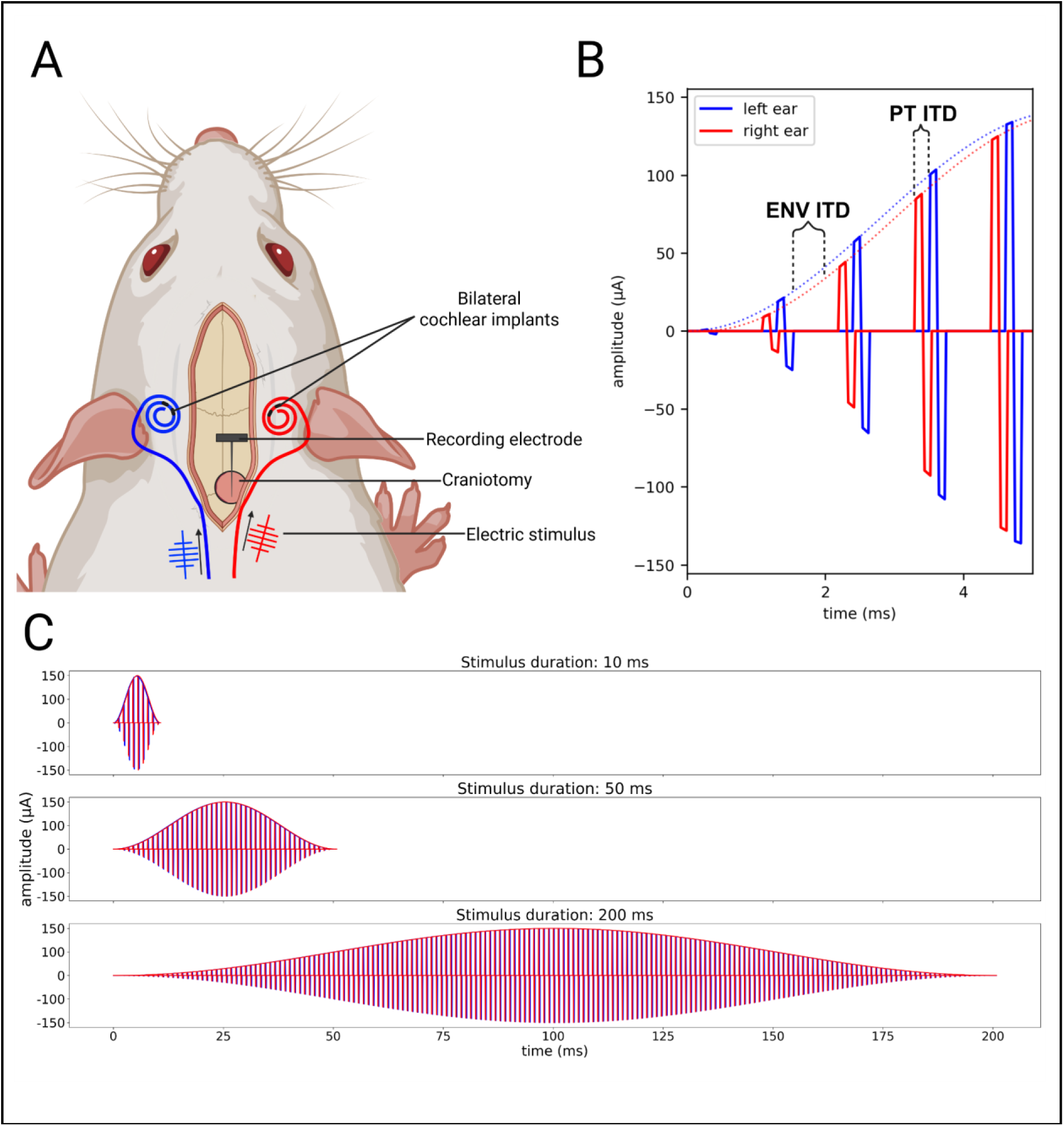
Illustration of stimulus. (A) Cartoon created with Biorender.com illustrating the experimental procedure. Rats were bilaterally implanted with cochlear implants via cochleostomy into the middle turn and a recording electrode array was inserted into the inferior colliculus (B) Hanning window envelopes were used to amplitude modulate biphasic pulse trains. Envelope (ENV) and the pulse timing (PT) ITDs could be varied independently of each other. In this example, the left ear (blue) ENV rises earlier than the right ear (red) ENV, illustrating a left leading ENV ITD. Meanwhile, the pulses in the left ear occur after the pulses in the right, illustrating a right leading PT ITD. (C) To address the possibility that ENV ITD sensitivity may depend on the steepness of the ENV, we tested three stimulus durations (10, 50, and 200 ms) with progressively sharper rising edges.

All electrical intracochlear stimuli consisted of pulse trains of biphasic, anode leading current pulses (duty cycle: 40.96 μs positive, 40.96 μs at zero, 40.96 μs negative). The generated binaural electrical pulse train contained ENV ITD and PT ITD, varying independently within {+80, 0, -80} µs, where negative values indicated that the left ear stimulus was leading. In total, there were 54 different stimulus combinations (3 ENV ITDs x 3 PT ITDs x 2 pulse rates x 3 modulation frequencies). We presented each of the 54 stimuli 15-30 times in a pseudo-randomized order.

### Data analysis

Data processing and analysis were performed using customized code written in Python. In total, 29 penetrations for the ND + acute CI cohort (n = 8), 25 penetrations for the NH + acute CI (n = 6), and 37 penetrations for the ND + chronic CI cohort (n = 6) were recorded.

#### Artifact rejection

CI stimulation pulses result in electrical artifacts characterized by sharp peaks in the recorded neural signal with a very high amplitude, typically much higher than peak values that are associated with normal neural spiking. In this study, electrical pulse train stimuli with durations of 10 ms, 50 ms and 200 ms were presented, resulting in an overlap between electrical artifacts and neural responses of IC neurons. A template subtraction approach was used to separate the artifacts from the neural responses caused by the stimulation (Hashimoto *et al*., 2002). The basic concept is to average across the repeats to generate the template and subtract the template from the individual signal. Given that linear subtraction preserves temporal properties of artifact residuals, we assumed the residuals, if any, would maintain temporal alignment with electrical pulses and appear consistently.

To gauge the success of the artifact rejection, we implemented a quality control procedure on the artifact-removed recording. First, we located the largest amplitude in the neural signal during CI stimulation. To ensure that individual large spike events were not incorrectly treated as artifacts, the amplitudes one interpulse interval before and after the maximum amplitude were extracted and summed to calculate a “triplet” value. The same procedure was performed on an equivalent post-stimulation recording time to determine the neural “baseline triplet”. Both triplet values were divided by the standard deviation of the last 100 samples of the recording, holding also a neural baseline without stimulation. The “triplets” during CI stimulation were compared to “baseline triplets” for all recordings of each channel. The recording was excluded for further analysis if “triplets” during CI stimulation exceed “baseline triplet” values by two times the 95th percentile. If more than ¼ of the presented repetitions of one stimulus combination were removed, the complete stimulus combination is excluded for the particular channel for further analysis.

#### Analog multi-unit activity

Analog multi-unit activity (AMUA) was computed from the recorded extracellular voltage traces as previously described in Kayser *et al*. (2007) and Schnupp *et al*. (2015). This method quantifies neural activity by analyzing the amplitude envelope of the electrode signal in the frequency range that contains action potentials (Kayser *et al*., 2007; Schnupp *et al*., 2015). To calculate the AMUA, the artifact rejected neural signals were first subjected to bandpass filtering using a 4th-order Butterworth filter with a frequency range of 0.3-6 kHz. The absolute value of the filtered signal was then obtained, and a subsequent lowpass filter was applied at 6 kHz. The response window was set to the same length as the stimulus duration, and the average amplitude of the AMUA signal during the stimulus period was computed to quantify the neural response to the stimuli. The resulting AMUA trace served as a measure of local multi-unit firing rates that is usually less noisy than multi-unit activity measures based on threshold crossings (see Fig. 1 in Schnupp *et al*., 2015).

#### ITD sensitivity analysis

To identify responsive multi-units in the IC, we first performed a Wilcoxon signed-rank test for the one stimulus combination with the smallest duration (duration = 10 ms, click rate = 900 pps, PT ITD = 80 µs and ENV ITD = - 80 µs), comparing the averaged AMUA amplitude during the stimulation time with the averaged baseline AMUA amplitude during the equivalent post-stimulus time window. Only multi-units showing significant differences (p < 0.05) were considered responsive and were included in subsequent analyses.

For each responsive multi-unit, we calculated the proportion of the trial-by-trail variance in multi-units response amplitude accounted for (explained by) changes in stimulus PT ITD or ENV ITD respectively, using an approach based on a traditional analysis of variance (ANOVA). The variance explained (VE) ratio was computed as the ratio of the sum of squares explained by each factor (V_factor_) to the total sum of squares (V_total_):

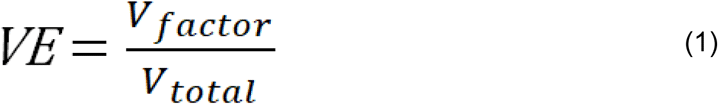

Where V_factor_ represents the sum of squares attributed to the factor, calculated as:

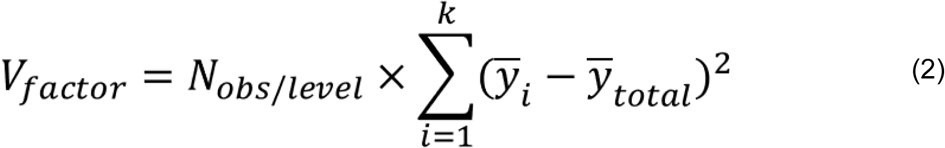

Here, N_obs/level_ is the number of repeats for a given factor “level”, *y*_i_ is the mean response at the i_th_ level, and *y*_total_ is the grand mean response. V_total_ was computed as the sum of squared deviations over all observations from *y*_total_. The analysis was conducted independently for the six combinations of click rates and durations (two click rates × three durations). Within each group, VE was calculated separately for PT ITD and ENV ITD effects.

A permutation test was used to decide whether VE values computed for each particular multi-unit were statistically significantly larger than zero. For each channel and condition, we performed 1000 iterations where all observations were randomly shuffled to create null distributions of VE values. The 95th percentile of these null distributions served as a significance threshold. Multi-units were therefore considered significantly sensitive to PT or ENV ITD respectively if the corresponding VE values exceeded their threshold.

## Results

Twenty biCI animals were used to investigate the neurophysiological sensitivity to PT ITD and ENV ITD in the IC using three different cohorts varying in hearing experience (Fig. 1). The total number of sampled multi-units across the three cohorts was: 1) 746 multi-units from AD + acute CI (n = 6). 2) 642 multi-units from ND + acute CI (n = 8). 3) 1073 multi-units from ND + chronic CI (n = 6). The total number of multi-units, as well as the number of artifact free multi-units confirmed by quality control included from each cohort for the different stimulus combinations, is given in Table 1 below.

**Table 1.**
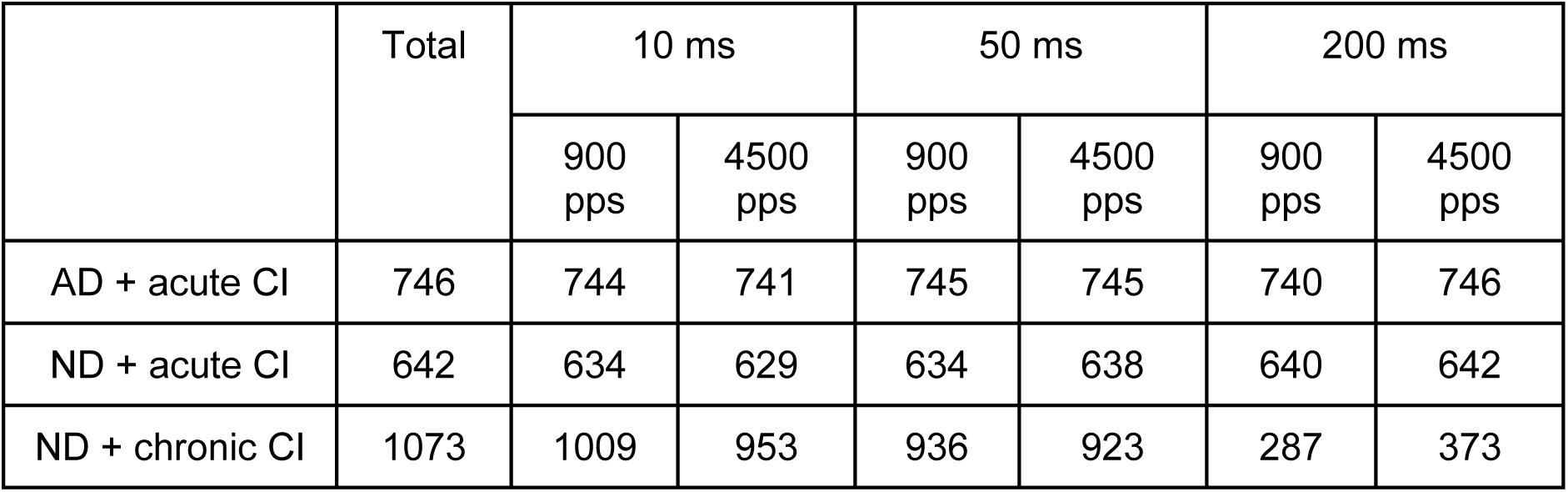
Number of used units for the analysis after artifact rejection and quality control for leftover artifacts.

### IC multi-unit activity in dependence of PT and ENV ITD

To calculate the overall neural strength of our signal, we used AMUA calculation as previously described in Kayser *et al*. (2007) and Schnupp *et al*. (2015). Figure 3 shows AMUA traces recorded from three illustrative example multi-units. The top row shows how responses vary if PT ITD is held constant at 0 µs and ENV ITD is varied (-80, 0, 80 µs), while the bottom row shows the converse conditions, with ENV ITD held constant at 0 µs while varying PT ITDs (- 80, 0, 80 µs). Displayed AMUA traces were averaged over all presented repetitions and normalized by the maximum amplitude for ease of comparison. *VE(PT)* decreases from left to right (Fig. 3, bottom row), while *VE(ENV)* remains minimal across all examples (Fig. 3, top row). The examples shown illustrate that the evoked responses change very little if ENV ITD changes, but the changes in PT ITD can induce clear changes in response shape and amplitude. Note that this figure does not include examples of multi-units exhibiting high *VE(ENV)* simply because our dataset contained no examples of multi-units with *VE(ENV)* values exceeding a few percent. We will examine the distributions of observed *VE(PT)* and *VE(ENV)* values further below. All multi-units exhibited onset responses, with some units (left and middle panels, blue traces) also showing secondary peaks at approximately 30 to 40 ms post-stimulus.

**Figure 3.**
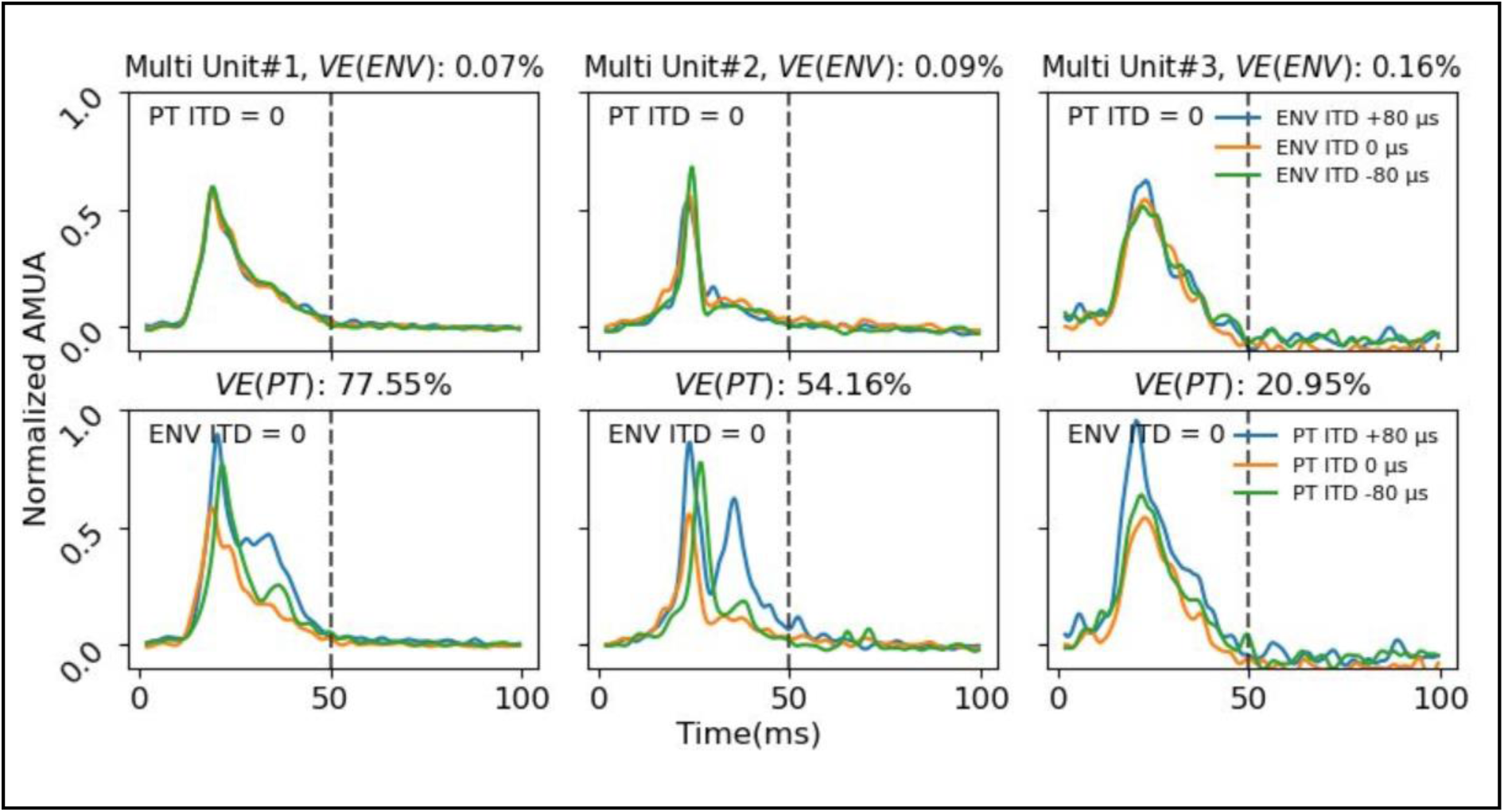
Sample of analog multi-unit activity (AMUA) responses from three different multi-units (columns) showing the effect of varying envelope ITD (ENV ITD) while holding PT ITD constant at 0 µs (top row) or varying pulse timing ITD (PT ITD) while holding ENV ITD constant at 0 µs (bottom row). The samples show responses to stimuli with a 50 ms duration (dotted vertical line). Each colored line shows the averaged evoked response over 30 stimulus presentations with the ENV ITD and PT ITD parameters indicated by the labels in the legends.

### Variance explained calculation exposes substantially greater sensitivity to PT ITD than to ENV ITD

The variance explained (VE) analysis quantified the relative contribution of PT ITD and ENV ITD to neural response variability in IC. Figure 4 shows scatter plots comparing the proportion of variance explained by PT ITDs versus ENV ITDs across different stimulus durations (10, 50, 200 ms) at 900 pps and for each experimental cohort. Each point in the scatter plot represents one multi-unit, and different color codes are used to represent different animals.

**Figure 4.**
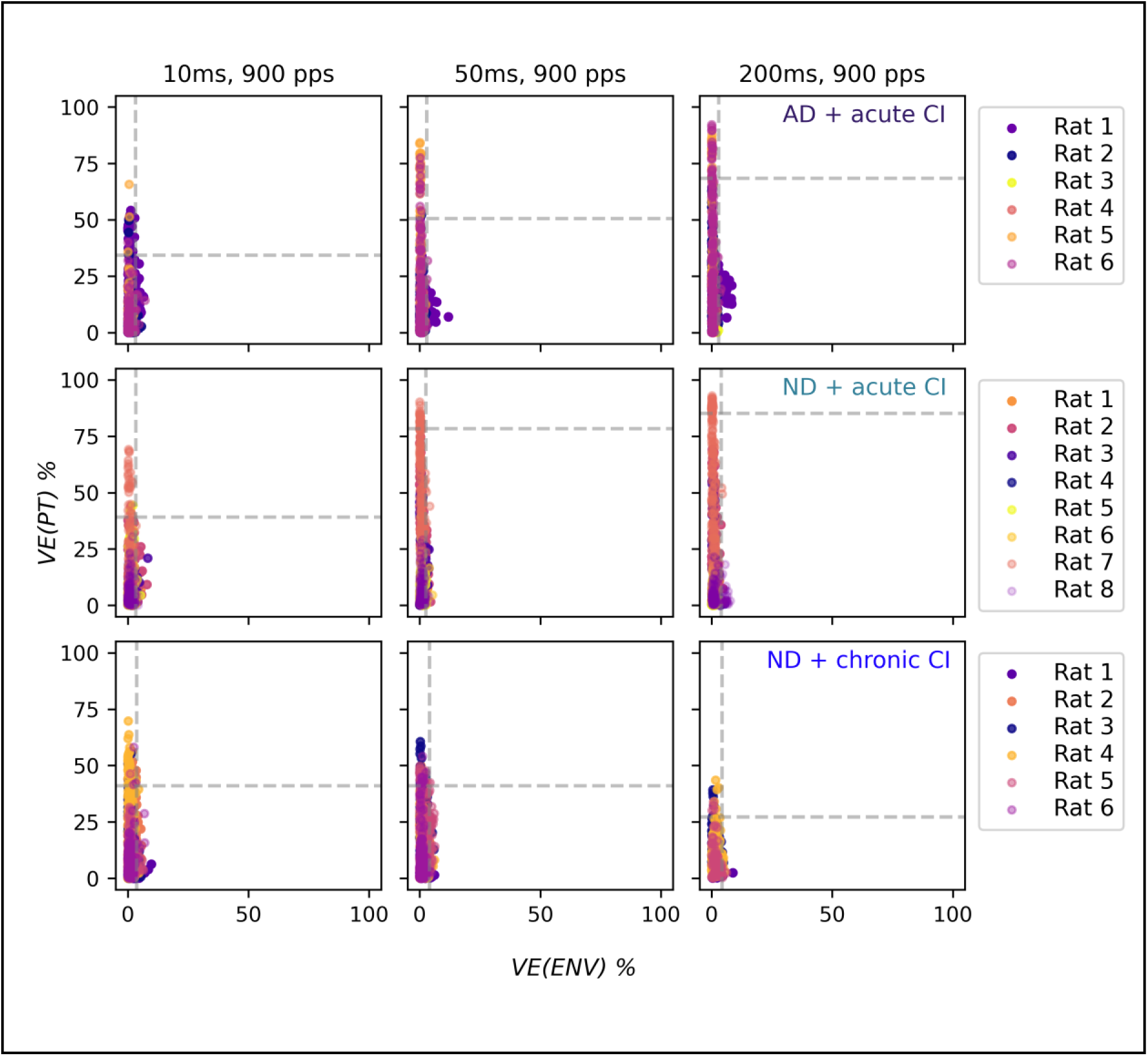
Increased variance explained (VE) values for pulse timing ITD (PT ITD) compared to envelope ITD (ENV ITD) for all three cohorts over all three durations (from left to right: 10 ms, 50 ms, 200 ms) at 900 pps. All dots from one animal are labeled with the same color. Upper row displays VE values for PT ITDs and ENV ITDs for the AD + acute CI cohort (n = 6). Middle row presents VE values calculated for the ND + acute CI cohort (n = 8). Lower row shows VE values for the ND + chronic CI cohort (n = 6). Values on the Y-axis indicate the percentage of VE explained by PT ITD. X-axis indicates the percentage of VE explained by ENV ITD. The dotted grey line indicates the 95th percentile of the data.

In all conditions, points spread over a wide range along the vertical *VE(PT)* axis, with sometimes over 80 % of the observed variance in responses attributable to PT ITD changes, demonstrating strong PT ITD tuning in many multi-units showed. In contrast, all points cluster close to zero relative to the *VE(ENV)* axis, with ENV ITD only rarely accounting for more than 10 % response variance, indicating that only a small part of the multi-units are weakly tuned to ENV ITD.

In the acute CI cohorts (AD + acute CI and ND + acute CI), as the stimulus duration increases, the 95th percentile of both *VE(PT)* and *VE(ENV)* increases. The 95th percentile of variance explained by PT ITD increases systematically with stimulus duration, rising from approximately 40 % at 10 ms to approximately 80 % at 200 ms. This trend is, however, not observed in the ND + chronic CI cohort, where the 95th percentile of *VE(PT)* decreases from approximately 40 % at 50 ms to approximately 30 % at 200 ms.

Although one might expect envelope features to be more salient in the shorter stimuli which have more rapidly rising envelopes, and that this might aid the interaural comparison of the timing of envelope features, no clear dependency of ENV ITD sensitivity on stimulus duration was observed in Figure 4. The *VE(ENV)* values remained constrained to 0–10 % across all conditions.

In addition to the 900 pps data shown in Figure 4, Figure 5 allows a direct comparison of the neurophysiological ITD sensitivity at stimulation rates of 900 and 4500 pps (indicated by the pale yellow background). VE values for ENV and PT ITD presentation at different stimulus durations are shown for each multi-unit, with the three experimental cohorts represented in separate columns. While the same general trends are observed as shown in Figure 4, the box plot representation provides a clear depiction of data distribution and facilitates the identification of outliers. Across all conditions, under the same stimulus condition, the median VE is consistently higher for PT than for ENV ITD cues, and the spreading of the *VE(PT)* is always wider than for *VE(ENV)*. Across all cohorts, a higher pulse rate (4500 pps) has kept approximately the same median value for *VE(PT)* across all stimulus durations. Accordingly, no improvement in envelope sensitivity was observed, in either the median values, the upper range, or the overall data distribution, with no notable changes across all conditions.

**Figure 5.**
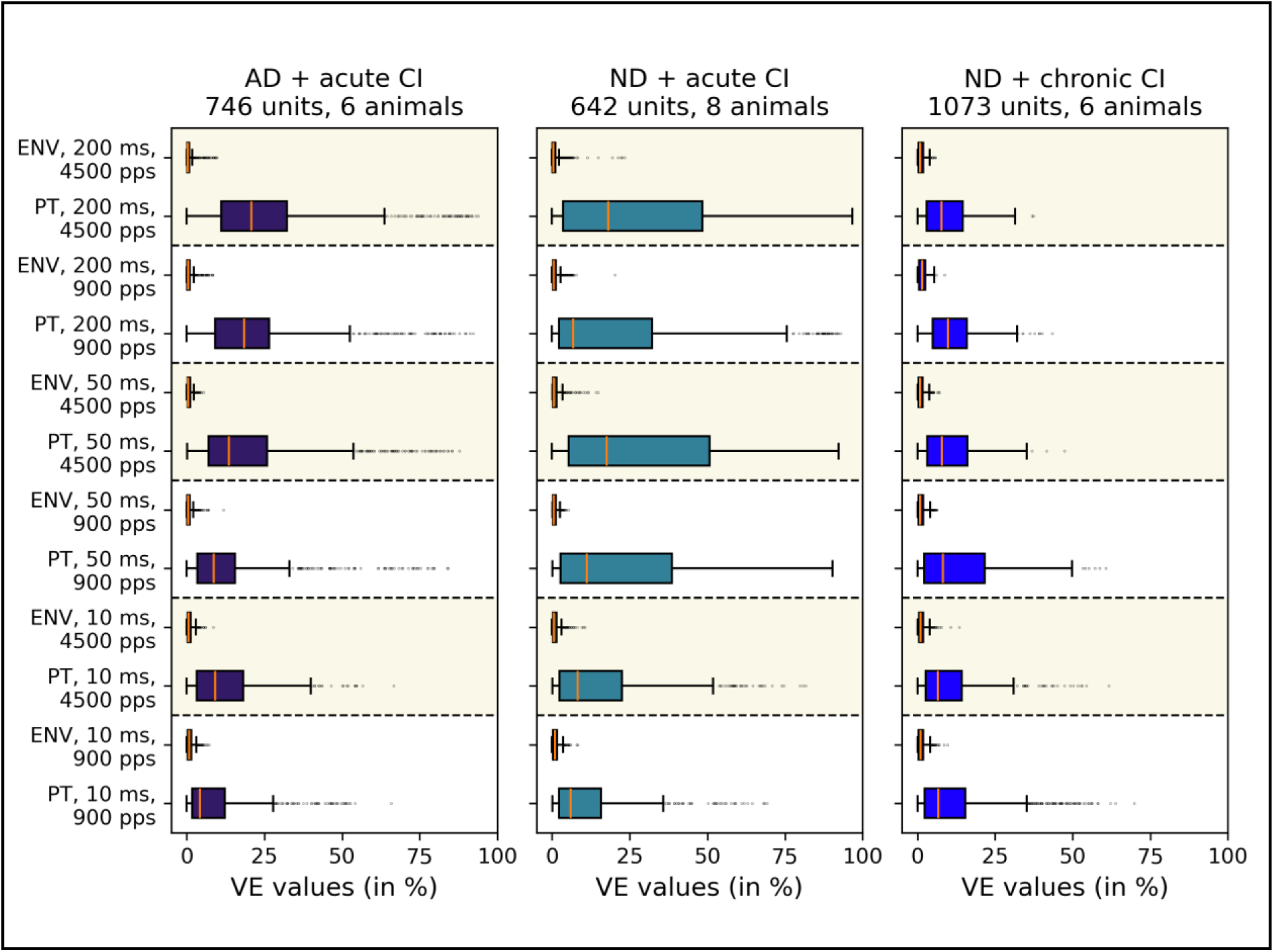
Direct comparison of variance explained (VE) values (in %) for pulse timing ITD (PT) and envelope ITD (ENV) for each unit displayed as box plots. Data are shown for three stimulus durations (10 ms, 50 ms, 200 ms; bottom to top) at 900 pps and 4500 pps stimulation rate (highlighted in pale yellow). Columns, as well as box colors, indicate the experimental cohorts (see subplot titles). Whiskers extend to 1.5× inter quartile range, and data points beyond this range are plotted as outliers.

### Neuronal PT ITD sensitivity dominates, while ENV ITD sensitivity is near chance level

Figure 6 shows the percentage of IC multi-units that exhibit VE values exceeding the 5 % significance threshold determined by the permutation test described in the Methods section. The proportion of IC units with responses significantly driven by PT ITD is consistently greater than those driven by ENV ITD for all tested stimulus parameters (Fig. 5). The proportion of PT ITD-sensitive multi-units exceeded 65 % across all cohorts and stimulus conditions. In contrast, the proportion of multi-units significantly sensitive for ENV ITD reached 10 % in only a single case (tested with 200 ms long stimuli at 900 pps), and generally hovered close to the expected 5 % false positive rate for this test. In other words, while PT ITD sensitivity was clearly widespread among IC multi-units, evidence for ENV ITD sensitivity was weak or absent. At 900 pps (Fig. 6, upper row), the proportion of PT ITD sensitive IC units increased with stimulus duration, with an exception in the ND + acute CI cohort, where the proportion decreased from 75.87 % to 68.91 % as duration increased from 50 to 200 ms. Interestingly, multi-units maintained high PT ITD sensitivity even at 4500 pps, as demonstrated in the lower row of Figure 6. Except for the ND + chronic CI cohort at 200 ms, a higher proportion of multi-units exhibited PT ITD sensitivity at 4500 pps. Although higher pulse rates theoretically allow better envelope reconstruction through increased sampling, the 4500 pps condition showed only minimal improvement in ENV ITD sensitivity, with the only notable increase observed in the ND + acute CI group at 50 ms (from 2.05 % to 7.21 %).

**Figure 6.**
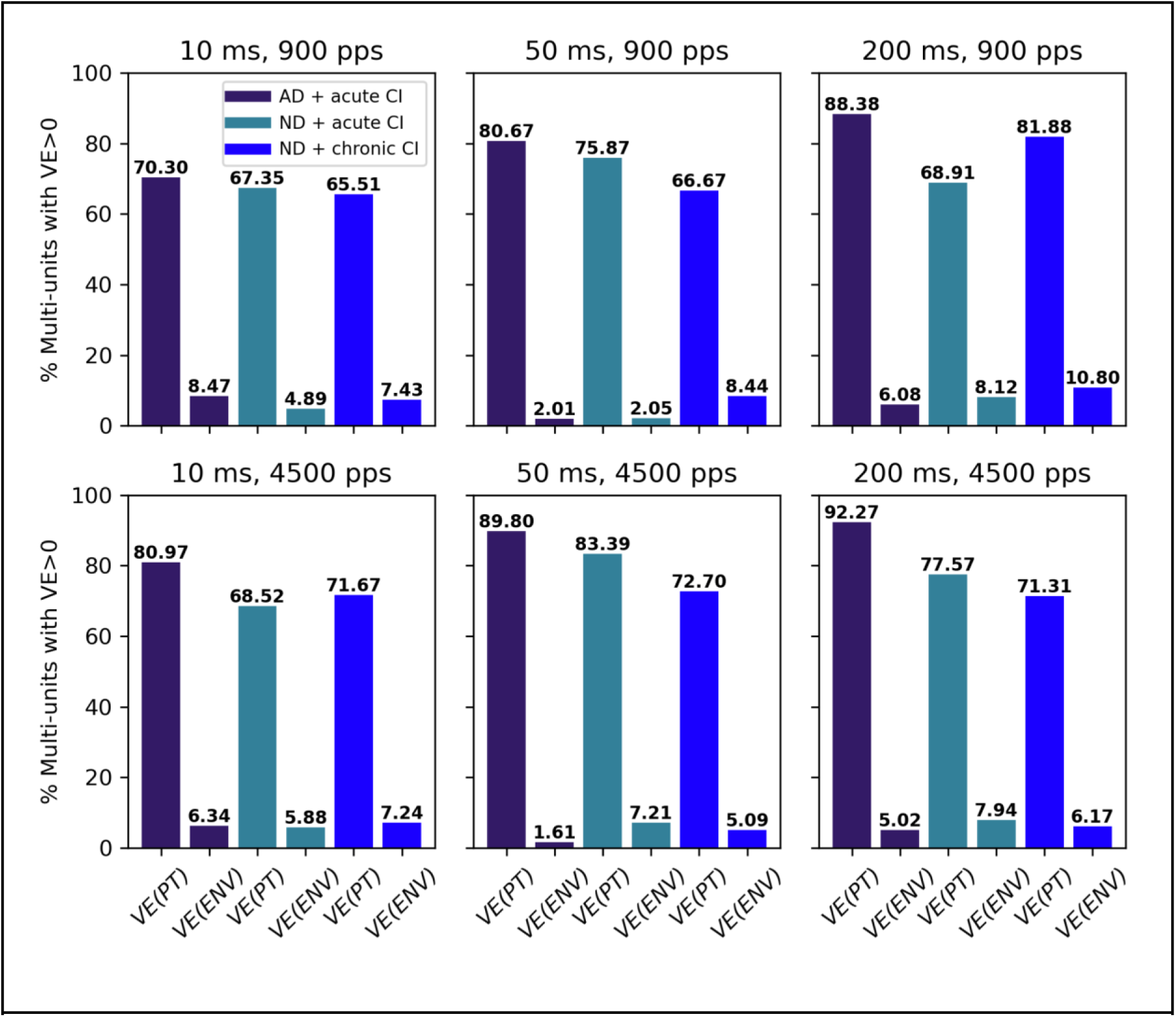
Proportion of multi-units that had VE values significantly greater than zero (y-axis) for the different cohorts and stimulus conditions. Significance was tested by permutation of VE values and testing if the corresponding unit exceeded 95 % confidence interval. Note that the proportion of multi-units with significant *VE(PT)* is much larger than that with significant *VE(ENV)* in all conditions. Top row: 900 pps stimuli, bottom row 4500 pps stimuli. Bar color indicates cohort (see legend).

## Discussion

The key finding of this study is that ITDs presented in the pulse timing of an electric stimulus elicited greater changes in neural activity in the IC than ITDs presented in the pulse train envelope. The CI stimuli we used evoked robust responses, but changing stimulus PT ITD routinely led to much greater changes in the neural responses than varying ENV ITD (compare example AMUA traces in Fig. 3). Quantifying these observations on the basis of VE calculation revealed that a high number of multi-units had significant, and often substantial, PT ITD sensitivity, while sensitivity to ENV ITD was comparatively very small or absent (Fig. 4 and 5). The percentage of units with VE values significantly greater than zero was about 8-12 times larger for *VE(PT)* than for *VE(ENV)*, regardless of stimulus type or hearing experience (Fig. 6). The timing of individual pulses in the CI stimulation therefore has a much larger effect on neuronal IC responses than the timing of pulse train envelopes, a fact largely ignored in current clinical CI processing strategies, with exception of the low frequency channels of the MED-EL FS4 strategy. Our results indicate that limited efforts made to encode auditory information in the timing of individual stimulus pulses in CI patients can result in substantially poorer sensitivity for ITD cues.

### Comparison to behavioral literature

In acoustic hearing, FS and ENV ITD cues are normally coupled to one another, given that ITDs will delay both FS and ENV features at the far ear by the same amount. Previous literature on ITD presentation on FS and ENV in normal hearing psychoacoustic experiments show much better FS than ENV ITD sensitivity using low-frequency AM tones, with FS thresholds between 60-180 μs and ENV thresholds often exceeding 1,000 μs (Moore *et al*., 2018). Bernstein & Trahiotis (2002) showed that ENV ITD sensitivity in NH listeners can be much better (as low as ∼100-130 μs) with high-frequency modulated or transposed stimuli. However, comparing ITD sensitivity directly between acoustic hearing (FS and ENV) and electrical hearing (PT and ENV) is problematic, due to many fundamental differences in the mechanism of natural cochlear sound transduction and electrical CI signal transduction, respectively. For example, in clinical CI stimulation, ENV and PT features are not coupled, and usually vary independently of one another, with ENV ITDs being informed by sound source location but PT ITDs mostly created by random temporal offsets between the left and right ear pulse train generators. Although previous literature on ENV and PT ITD presentation in CI patients by Noel & Eddington (2013), Majdak *et al*. (2006), and Laback *et al*. (2004) found PT ITD sensitivity when testing at lower pulse rates, the ITD thresholds for PT ITD presentation at high pulse rate (around 900 pps) were high. These findings are in contrast to various behavioral studies from our groups, like Buck *et al*. (2023) or Schnupp *et al*. (2025), where we demonstrated robust behavioral sensitivity in biCI rats to even quite small ITDs, and even at relatively high pulse rates of 900 pps. However, it is reasonable to assume that prelingually deaf biCI users, who are exposed to confusing or uninformative PT ITDs over many months or even years, may lose sensitivity to these cues. While the patients studied by Noel & Eddington (2013), Majdak *et al*. (2006), and Laback *et al*. (2004) were quite heterogeneous with respect to their clinical history, prior normal hearing experience, or the duration of hearing experience with clinical processors delivering misleading PT ITD cues, our cohorts differed only with respect to whether or not they had normal hearing experience in early life or electric hearing experience, and we did not find that these variables had a big impact on the observed neural tuning properties, as all cohorts showed much more sensitivity to PT ITD than ENV ITD (Fig. 4 and 5) and the proportion of multi-units with *VE* > 0 was similar in all cohorts (Fig. 6). The only apparent difference between cohorts in our dataset is that the 95th percentile of *VE(PT)* decreased with increasing stimulus duration in the ND + chronic CI cohort, but it remained stable or tended to increase in the other two cohorts (Fig. 4 and 5). However, given the relatively small cohort sizes, the high within-cohort variability, and the lack of statistical independence between data points recorded from any one array electrode penetration or animal, we cannot exclude the possibility that this apparent difference between cohorts may simply reflect a sampling effect. Multi-units with either large or small *VE(PT)* values had a strong tendency to cluster anatomically, and how long the “tail” of the *VE(PT)* distribution sampled from any one animal or cohort depends in large part on how often the multi-electrode penetrations happened to strike clusters of neurons with large *VE(PT)*.

All early deafened biCI animals of Schnupp *et al*. (2025) had shown that PT ITDs dominate their lateralization decision when tested with stimuli that are essentially identical to the ones used in the electrophysiological recordings here. Our electrophysiological observations are thus in excellent agreement to this behavioral result and suggest that PT ITD very strongly modulates both IC neural responses and psychoacoustic lateralization judgments, while ENV ITD does so only very weakly. By investigating additional cohorts in this paper, which differed greatly in their hearing experience prior to electrophysiological recordings (one neonatally deafened with essentially no hearing experience prior to recording, another with normal hearing experience until the time of recording), we can now conclude that the dominant role of PT over ENV cues is a robust phenomenon that is not strongly influenced by auditory input during early development.

### Comparison to previous electrophysiological work in CI animals

Our results are perhaps unsurprising when considered in the light of a previous study by Hancock *et al*. (2017), who studied responses of midbrain neurons to amplitude modulated pulse trains delivered via CI, and who observed that the IC neuron responses were frequently locked to the timing of suprathreshold pulses (shown in Fig. 8 in Hancock *et al*., 2017). It is to be expected that the timing of spikes of brainstem neurons that feed into the binaural comparison circuits in the superior olivary nuclei will similarly synchronize to the timing of individual stimulus pulses, rather than to temporal features of the pulse train ENV. Another study by Dynes & Delgutte (1992) also confirms this theory, as they found that electrical stimulation can produce significant phase locking of the auditory nerve well above 1000 pps.

Unlike the very sharp flanks of individual suprathreshold pulses, which can directly trigger action potentials in auditory nerve fibers, the ENV cannot be directly observed by the auditory nerve or brainstem, rather, it must be inferred (“reconstructed”). Consequently, we would only expect ENV ITD to be an effective binaural cue in CI stimulation if, or insofar as, mechanisms early in the auditory brainstem are able to reconstruct temporal features of the envelope of a stimulus pulse train with a higher temporal resolution than the pulse rate. In the absence of such an ENV feature reconstruction either in the cochlear nucleus or in the nuclei of the superior olive, the timing of ENV features would inevitably be “rounded up” to the timing of the next electric pulse triggered action potential, and PT ITDs would effectively “overwrite” ITDs of ENV features. The clear dominance of PT ITDs over ENV ITDs documented here, which occurs at all tested parameters and all different hearing experience conditions tested, can thus be explained if we simply assume that there is only little or no “envelope reconstruction filtering” in the early stages of the CI stimulated auditory pathway, up to and including the superior olivary nuclei. Note that this situation is fundamentally different from that of normal, hair-cell based, acoustic hearing, given that the cell membrane of inner hair cells can act like an envelope reconstruction filter for high frequency sound waves (Palmer & Russell, 1986). In a normally hearing auditory nerve, action potential distributions can therefore track temporal ENV features of high frequency sounds smoothly and continuously, unlike in a CI stimulated auditory nerve, where the timing of action potentials is “forced” onto the “temporal grid” laid down by the timing of the electrical pulses. These fundamental differences in temporal feature encoding imply that studies of ENV ITD sensitivity in normal, acoustic hearing can be poor predictors for what to expect in the CI stimulated case.

For example, Greenberg *et al*. (2017) have shown in guinea pig IC that acoustic stimuli with rapid (“ramped”) ENV onsets elicited much stronger ITD sensitivity than ones with slow onsets (“ramped”). The best neural just noticeable difference (JND) for a steep “ramped” envelope was ∼91 μs versus ∼407 μs for a slow “ramping” ENV. This is consistent with psychophysical data from Noel & Eddington (2013) as well as Klein-Hennig *et al*. (2011) showing that fast onsets and long interpulse phases result in lowest ITD thresholds. The main take-away from these studies is that the auditory system is most sensitive to ENV ITDs when the ENV has sharp rises and well-separated pulses (Klein-Hennig *et al*., 2011; Noel & Eddington, 2013; Greenberg *et al*., 2017). In contrast, our results show that, for pulsatile electric stimuli delivered at a range of different pulse rates, ENV ITD sensitivity remains uniformly poor irrespective of the slope of ENV onsets, as we would expect if we assume that the “rounding to the timing of the first supra-threshold pulse” that we just described is indeed a major limitation of temporal processing of CI pulse train stimuli.

Previous studies on congenitally deaf cats with biCIs or CI-stimulated rabbits have reported relatively poor ITD tuning for IC neurons tested with ITDs at high pulse rates (Hancock *et al*., 2010; Day *et al*., 2012; Chung *et al*., 2016). However, a number of design choices, including use of coarse ITD sampling with the large majority of ITDs sampled outside the physiological range, or the exclusion of onset responses in the study design, may have led these studies to underestimate the ITD sensitivity that may be achievable with CI stimulation under more optimal conditions, and we have previously reported that IC neurons of ND biCI rats that were tested predominantly in the physiological range exhibited ample innate sensitivity even to very small ITDs (Buck *et al*., 2021).

The first studies to show this remarkably good sensitivity to CI ITD stimuli even in the absence of hearing experience (Buck *et al*. 2021, Rosskothen-Kuhl *et al*. 2021, Buck et al 2023) used stimuli where ENV and PT ITDs covaried, as they normally would in acoustic hearing. However, this experimental design does not shed any light on whether neural ITD sensitivity is triggered by one of these or a combination of both. The results presented here now provide a clear answer to this question in favor of PT ITDs. The only animal biCI study known to the authors that precisely discriminates between ITD sensitivity for PT and ENV ITD was done by Smith & Delgutte (2008). Similarly to our AD + acute CI cohort, cats were deafened in adulthood, and 7-14 days later, acutely implanted on both sides prior to electrophysiological single-unit recordings. Smith & Delgutte (2008) presented 1000 pps and 5000 pps pulse trains with fixed PT ITD but continuously, slowly varying ENV ITD. Smith and Delgutte concluded that many IC neurons were sensitive to ITD presentation in ENV and PT for respective frequencies. By comparing the effect of ENV ITD modulation to that of changing PT ITDs in fixed step sizes, they observed a higher sensitivity to small steps in PT ITD than in ENV ITD at high pulse rates typically used in clinical settings (1000 pps). In that regard, our results in CI rats of different hearing experiences are in generally good agreement with those by Smith & Delgutte (2008). One difference we note, however, is that Smith & Delgutte (2008) observed almost no neural sensitivity to FS ITDs at 5000 pps, while we still observe many multi-units with significant PT ITD tuning at 4500 pps (Fig. 6 bottom). At present we do not know whether these differences are due to species differences or methodological differences. In any event, our results support Smith and Delgutte’s overall conclusion that a coding strategy of biCIs should control precise timing of current pulses based on the fine timing of the sound source in order to convey ITD cues successfully.

Finally, we would like to draw attention to two publications that corroborate our findings that multi-units in the IC display greater PT ITD sensitivity than ENV ITD sensitivity. Firstly, studies by Thompson *et al*. (2021) in ND biCI cats showed that using a research processor with a customized “ITD-aware” stimulation strategy, thereby presenting informative PT ITDs, results in better neurophysiological ITD sensitivity compared to using CIS-based coding strategies. Secondly, Sunwoo *et al*. (2021) stimulated ND biCI rabbits with the experimental “fundamental asynchronous stimulus timing” (FAST) coding strategy, where PT is extracted from the temporal features of incoming sounds and thereby informative. They found that experience with FAST improved ITD sensitivity in the IC compared to animals with no stimulation experience.

## Conclusion

In normal hearing, ITDs serve as powerful cues for determining the source of sound. Our research demonstrated that small ITDs are only neurophysiologically relevant under biCI stimulation if they are transmitted via pulse timing rather than the envelope of the stimulus, regardless of the envelope shape, the pulse duration, the pulse rate, or the hearing experience. The inadvertent and frequently contradictory PT ITDs generated by the majority of existing clinical processors using CIS-based coding strategies likely prevent the auditory system from accessing potentially valuable binaural information. With few exceptions (notably low frequency channels of the MED-EL FS4 strategy), current clinical devices do not encode the precise temporal details of acoustic signals with the sub-millisecond accuracy required. Our results suggest that this absence of precisely timed stimulation pulses in contemporary clinical processing is likely a major contributing factor for suboptimal binaural hearing performance seen in biCI patients.

## Additional information section

### Data availability statement

All data supporting the findings of this study with n < 30 are fully included in the manuscript figures. All data (n > 30) as well as the analysis code used to generate all the figures and statistical results included in this manuscript can be downloaded from https://auditoryneuroscience.org/dataShare.

### Author contributions

**S. Fang:** Conceptualization, Data curation, Software, Validation, formal analysis, Writing original draft, Writing – review & editing. **T. Fleiner:** Conceptualization, Data curation, Software, formal analysis, Validation, Writing original draft, Writing – review & editing. **F. Peng:** Data curation, Software. **S**. **Buchholz:** Data curation. **M. Zeeshan:** Data curation. **N. Rosskothen-Kuhl:** Conceptualization, Resources, Supervision, Validation, formal analysis, Funding acquisition, Writing original draft, Writing – review & editing. **J.W. Schnupp:** Conceptualization, Resources, Software, Supervision, Validation, formal analysis, Funding acquisition, Writing original draft, Writing – review & editing.

## Acknowledgements

We thank B. Castellaro for surgical support. We thank E. Becker for support with training the animals.

